# Radiosynthesis and Evaluation of Novel Cholesterol 24-Hydroxylase Positron Emission Tomography Tracers

**DOI:** 10.64898/2026.01.25.701607

**Authors:** Yinlong Li, Haofeng Shi, Zhendong Song, Taoqian Zhao, Yanwei Jiang, Danielle E. Hoyle, Jiahui Chen, Xin Zhou, Qilong Hu, Xiaoyan Li, Lingxin Meng, Ruihu Song, Zhenkun Sun, Achi Haider, Hongjie Yuan, Steven H. Liang

**Affiliations:** Department of Radiology and Imaging Sciences, Emory University, 1364 Clifton Road, Atlanta, Georgia 30322, United States; Department of Pharmacology and Chemical Biology, Emory University School of Medicine, Atlanta, Georgia, 30322, United States; Wallace H. Coulter Department of Biomedical Engineering, Georgia Institute of Technology and Emory University, Atlanta, Georgia 30332, United States

**Author notes:** Corresponding authors: Y. Li; S.H. Liang. These authors contributed equally (Yinlong Li, Haofeng Shi).

**Keywords:** Cholesterol 24-hydroxylase, Radiotracer, [^18^F]Cholestify, Positron emission tomography, Autoradiography, Structure–activity relationship

## Abstract

Cholesterol 24-hydroxylase (CH24H or CYP46A1) is a pivotal enzyme in brain cholesterol metabolism and has emerged as a therapeutic and imaging target in neurodegenerative disorders. Although [^18^F]Cholestify ([^18^F]CHL-2205) has shown promise as a positron emission tomography (PET) tracer for imaging of CYP46A1, the impact of cyclopropyl moiety conformation on binding and imaging performance remains unexplored. Here, we report the rational design and preliminary evaluation of novel CYP46A1 PET tracers, in which the left-side cyclopropyl group was modified into bridged, spirocyclic, and fused bicyclic architectures to probe steric and conformational effects. All compounds **9**–**11** exhibited high CYP46A1 affinity (IC_50_ = 0.19–0.28 nM). Radiosynthesis of [^18^F]**9**–**11** was achieved *via* copper-mediated [^18^F]fluorination, providing practical non-decay-corrected radiochemical yields of 10–34% with excellent radiochemical purity (>98%). *In vitro* autoradiography in rat brain sections demonstrated specific and regionally selective binding, comparable to that observed for [^18^F]CHL-2205. These cyclopropyl-derived scaffolds establish a scaffold-driven strategy for PET tracer development, providing a robust framework for further structure–activity relationship studies and the rational optimization of CYP46A1 PET tracers.

## INTRODUCTION

The central nervous system (CNS) contains approximately 23% of the body’s total cholesterol, and tight regulation of cholesterol homeostasis is essential for neuronal function and neural circuit integrity.^1-4^ In contrast to peripheral tissues, CNS cholesterol metabolism is largely isolated from systemic circulation by the blood–brain barrier (BBB).^5^ Oxidative cholesterol metabolism is mediated by members of the cytochrome P450 (CYP450) superfamily, including the CYP7, CYP27, and CYP46 subfamilies in mammals.^6^ Among these enzymes, cholesterol 24-hydroxylase (CYP46A1, also known as CH24H) is uniquely expressed in the CNS and represents the principal CYP450 enzyme responsible for cerebral cholesterol turnover. CYP46A1 is predominantly localized in neurons within cognitive- and motor-related brain regions, including the cerebral cortex, hippocampus, and striatum.^7^ By catalyzing the conversion of cholesterol to 24S-hydroxycholesterol (24S-OHC), CYP46A1 mediates the primary pathway for cholesterol elimination from the CNS. ^8^ This more polar metabolite readily crosses the BBB into the systemic circulation for hepatic clearance, thereby preventing pathological cholesterol accumulation and preserving cerebral cholesterol homeostasis.^7, 9^ Consistent with this central role, dysregulation of CYP46A1 has been strongly implicated in the pathogenesis of multiple neurodegenerative disorders, including Alzheimer’s disease (AD), Huntington’s disease (HD), and Parkinson’s disease (PD).^10, 11^ In AD-related studies, upregulation of brain CYP46A1 enhances cholesterol clearance, which in turn reduces amyloid-β (Aβ) production and deposition, thereby alleviating neuropathology.^12^ At physiological concentrations, the CYP46A1-derived metabolite 24S-OHC also acts as an endogenous ligand for neuronal liver X receptors (LXRs), activating downstream signaling pathways that confer neuroprotective effects and attenuate Aβ-associated neuroinflammation.^13, 14^ Restoration of CYP46A1 activity in HD models has been shown to normalize sterol homeostasis and significantly improve motor performance.^15^ In contrast, increased CYP46A1 expression has been observed in PD, which has been associated with enhanced α-synuclein pathology and dopaminergic neurodegeneration.^16^ These findings underscore the disease-specific role of CYP46A1 dysregulation in neurodegenerative disorders and highlight the need for tools to quantitatively assess CYP46A1 expression and function *in vivo*.^9^

Positron emission tomography (PET) is a powerful molecular imaging modality that enables noninvasive, highly sensitive quantification and visualization of biological processes *in vivo*.^17-20^ PET imaging of CYP46A1 offers a unique opportunity to monitor enzyme activity in the brain, providing a valuable tool for early detection and disease progression assessment. As shown in **Figure 1A**, the first-generation tracer [^11^C]**1** exhibited moderate affinity and limited brain uptake, restricting its application to preclinical studies.^21^ Subsequent tracers [^18^F]**3g** (IC_50_ = 8.7 nM) and [^18^F]T□008 (IC_50_ = 8.8 nM) demonstrated markedly improved affinity and preferential accumulation in CYP46A1-rich regions nonhuman primates (NHPs).^22, 23^ More recently, the structurally distinct [^18^F]CHL-2310 was validated for specific binding and favorable kinetic properties.^24, 25^ Notably, [^18^F]Cholestify ([^18^F]CHL-2205) has been extensively assessed through cross-species postmortem analyses in rodents, nonhuman primates, and humans, confirming high affinity and selectivity for CYP46A1.^26^ A deuterated analog, [^18^F]CHL□2205□*d*_*3*,_ was also developed, showing comparable binding and imaging performance.^27^ Building on the [^18^F]CHL-2205 scaffold, we report the modification of the left-side cyclopropyl group into bridged, spirocyclic, and fused bicyclic architectures as a rational strategy to interrogate the roles of steric hindrance and conformational rigidity in CYP46A1 PET tracer design (**Figure 1B**).

**Figure 1.**
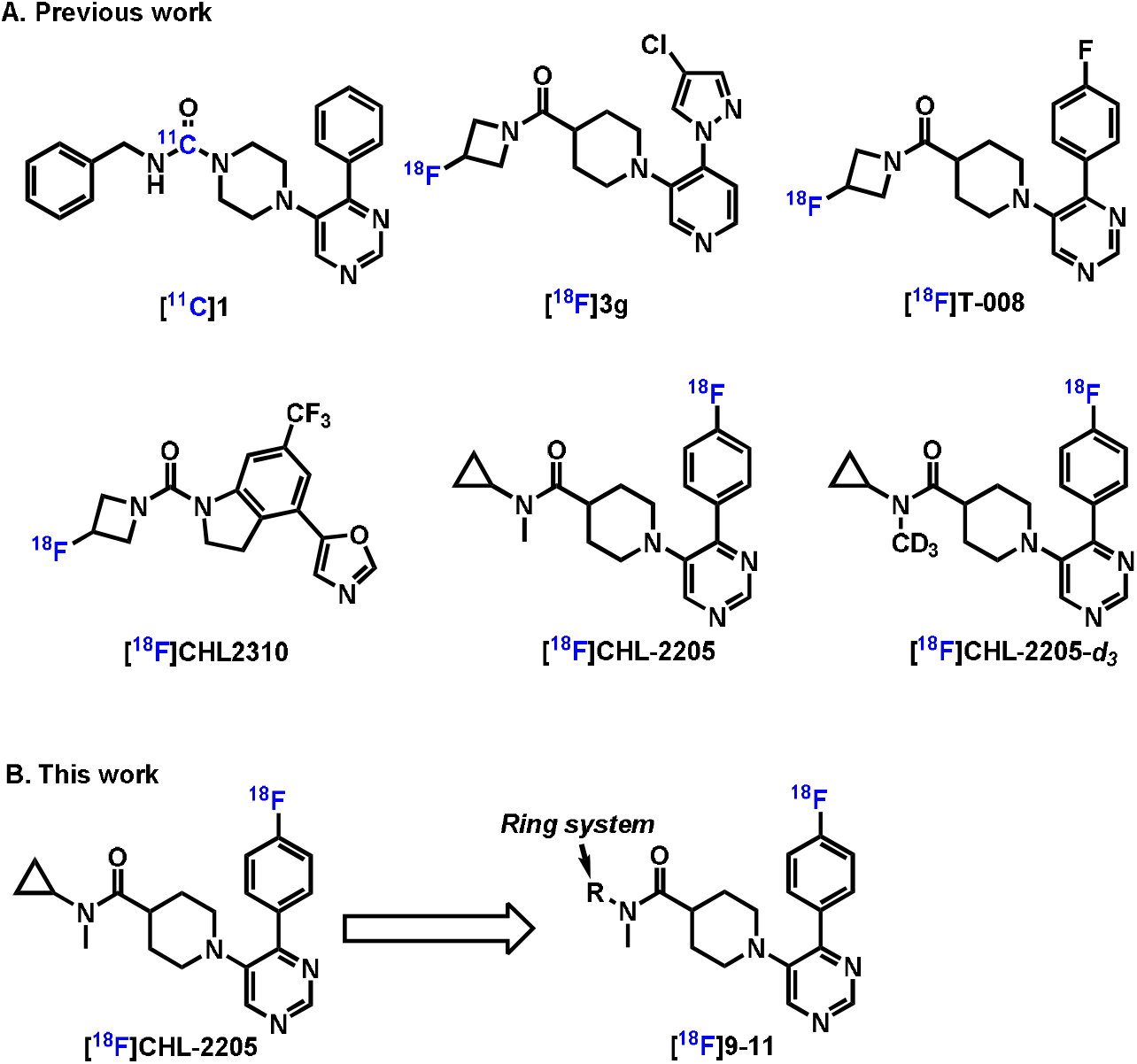
(A) Reported CYP46A1 PET tracers; (B) this study.

## RESULT AND DISCUSSION

### Chemistry

Compounds **8**–**10** were synthesized by coupling 1-(4-(4-fluorophenyl)pyrimidin-5-yl)piperidine-4-carboxylic acid (**2**) with the corresponding amines (**3**–**5**) using HATU and DIEA in DMF, affording the intermediate amides **6**–**8** in good yields (≥67%). Subsequent *N*-methylation with NaH and MeI in THF provided the target compounds **9**–**11** in 49– 70% yields (**Scheme 1**).

**Scheme 1.**
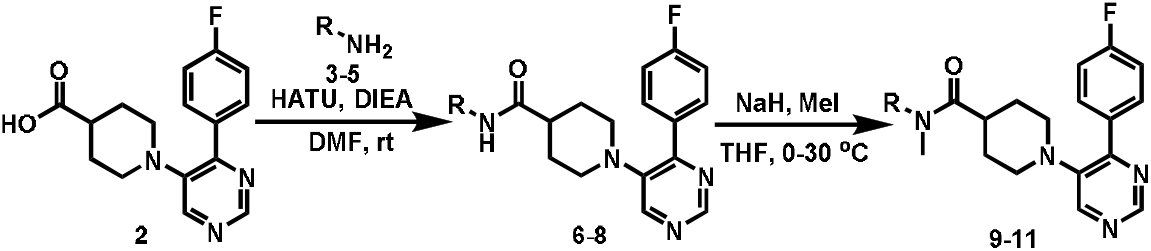
Synthesis of compounds **9-11**.

### Pharmacology

As illustrated in **Table 1**, compounds **9**–**11** exhibited high inhibitory potency toward CYP46A1, with IC_50_ values in the low-nanomolar range (0.19–0.28 nM), as determined by the radioligand binding assay (RBA). All three analogs demonstrated potency comparable to compound CHL-2205 (IC_50_ = 0.27 nM), indicating that modification of the left-side cyclopropyl group does not compromise CYP46A1 target engagement. From a physicochemical perspective, compounds **9**–**11** displayed favorable properties for CNS PET tracer development, with moderate lipophilicity (logP = 2.81–3.06; logD = 3.44–3.66), low polar surface area (tPSA = 49.33 Å^2^), and absence of hydrogen bond donors (HBD = 0). These parameters are consistent with favorable BBB permeability and align well with the predicted logBB values (−0.12 to 0.27). All three compounds also achieved MPO scores above 5, further supporting a balanced optimization of potency and CNS-relevant properties.

**Table 1.** *In vitro* CYP46A1 inhibitory potency and physicochemical properties of compounds **9**–**11**. ^*a*^Values were calculated with ChemDraw 21.0 software. ^*b*^Values were predicted with ACD/labs. ^*c*^Determined by “shake flask” method.

| Compound# | mw<sup>a</sup> | IC<sub>50</sub> (nM) | logP<sup>b</sup> | logD<sup>c</sup> | tPSA<sup>b</sup> | HBD<sup>b</sup> | logBB<sup>b</sup> | MPO score |
| --- | --- | --- | --- | --- | --- | --- | --- | --- |
| 9 | 380.5 | 0.19 | 2.81 | 3.44 | 49.33 | 0 | -0.12 | 5.5 |
| 10 | 394.5 | 0.28 | 2.93 | 3.47 | 49.33 | 0 | 0.27 | 5.3 |
| 11 | 394.5 | 0.24 | 3.06 | 3.66 | 49.33 | 0 | -0.04 | 5.2 |

### Radiochemistry

Based on the radiolabeling strategy previously reported for CHL2205, the boronic pinacol ester (Bpin) derivatives **19**–**21** were designed as the radiolabeling precursors (**Scheme 2A**). Briefly, carboxylic acid **12** was coupled with the corresponding amines (**3**–**5**) using HATU and DIEA in DMF to yield amide intermediates **13**–**15** in good yields (55–96%). Subsequent *N*-methylation with NaH and methyl iodide in THF provided amides **16**–**18** in 59–88% yields. Introduction of the Bpin moiety was carried out *via* palladium-catalyzed Miyaura borylation to deliver the desired precursors **19**–**21** in moderate yields (31–57%). As shown in **Scheme 2B**, The radiosynthesis of [^18^F]**9**–**11** was achieved through Cu(py)_4_(OTf)_2_ mediated radiofluorination in the presence of tetraethylammonium bicarbonate (TEAB) in a *N,N*-dimethylacetamide (DMAC)/n-BuOH solvent system at 110 °C for 15 min. [^18^F]**9**–**11** were obtained in 10–34% non-decay-corrected radiochemical yields, with excellent radiochemical purity (>99%) and high molar activity (>35 GBq/μmol). The *in vitro* stability of these tracers was subsequently evaluated in phosphate-buffered saline (PBS). All tracers remained intact for at least 90 min, indicating excellent stability and warranting further biological evaluation (**Figure 2**).

**Scheme 2.**
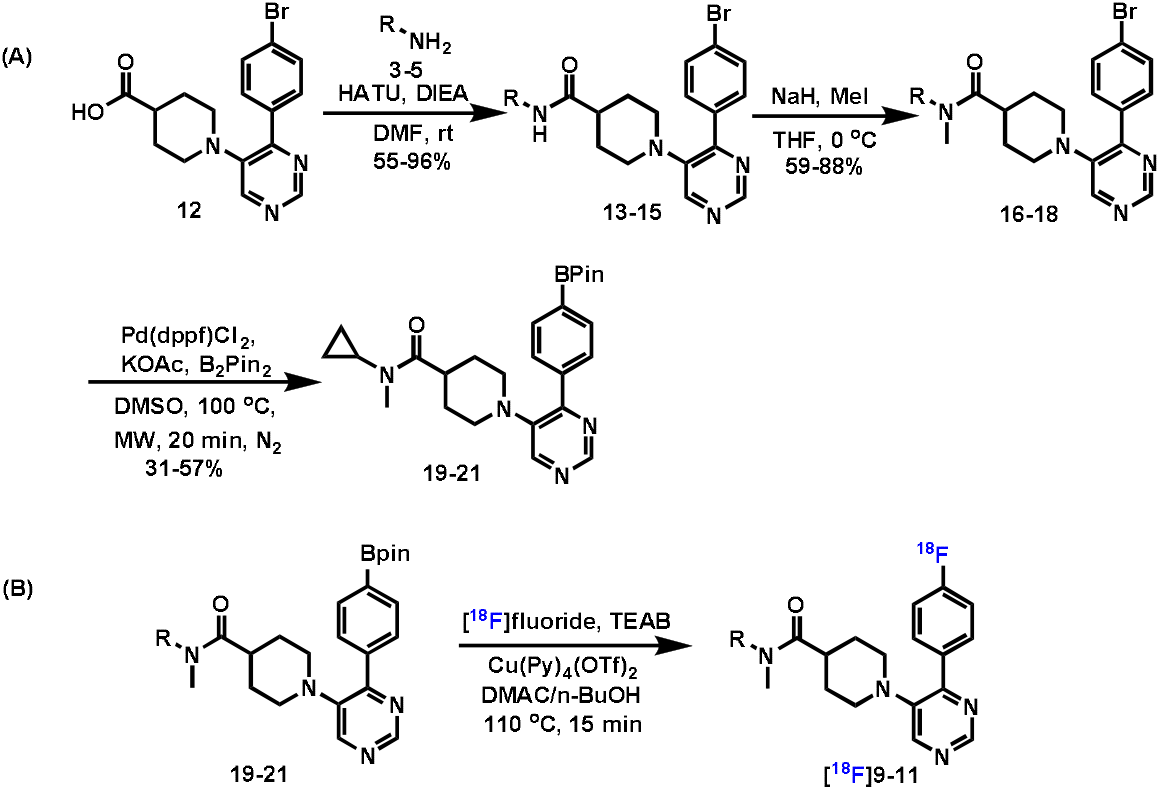
(A) Synthesis of precursor **19-21**. (B) Radiosynthesis of [^18^F]**9-11**.

**Figure 2.**
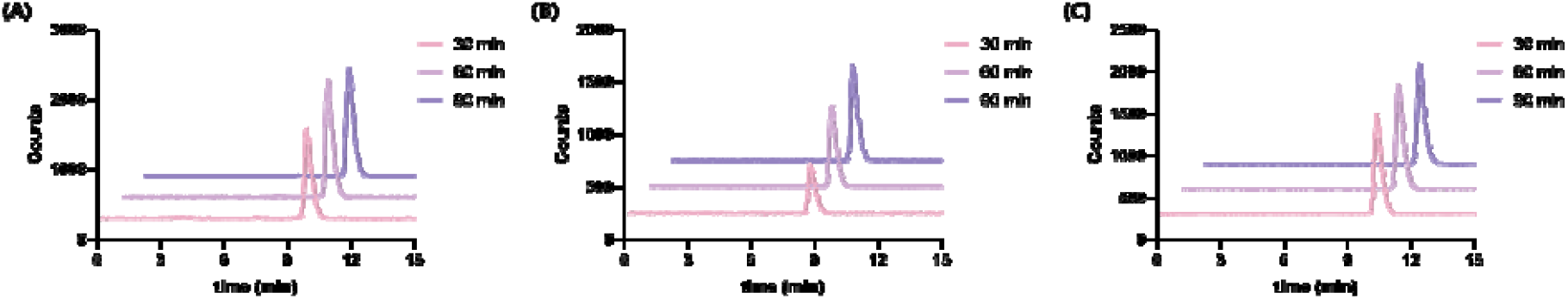
*In vitro* stability study of (A) [^18^F]**9**, (B) [^18^F]**10**, (C) [^18^F]**11** in PBS.

### *In Vitro* Autoradiography Study

To assess the regional distribution and binding selectivity of [^18^F]**9**–**11**, *in vitro* autoradiography studies were performed using sagittal rat brain sections (**Figure 3**). All three tracers displayed a heterogeneous and regionally selective distribution pattern, characterized by high radiotracer accumulation in the thalamus, striatum, cortex, and hippocampus, whereas the cerebellum showed consistently low signal intensity. This regional uptake pattern is consistent with the reported expression profile of CYP46A1 in the brain.^28, 29^ Notably, the overall binding patterns of [^18^F]**9**–**11** were comparable to those previously reported for [^18^F]CHL-2205, indicating that modification of the left-side cyclopropyl moiety into bridged, spirocyclic, or fused bicyclic architectures does not disrupt CYP46A1 target engagement at the tissue level. To evaluate binding selectivity, brain sections were co-incubated with an excess of the corresponding unlabeled reference compounds (**9**–**11**) or the selective CYP46A1 inhibitor soticlestat. Under these conditions, radiotracer binding was markedly reduced across all examined brain regions, confirming that the observed signals are predominantly attributable to specific binding to CYP46A1. These results demonstrate that [^18^F]**9**–**11** retain high specificity and appropriate regional distribution, supporting their further development as CYP46A1-targeted PET imaging probes.

**Figure 3.**
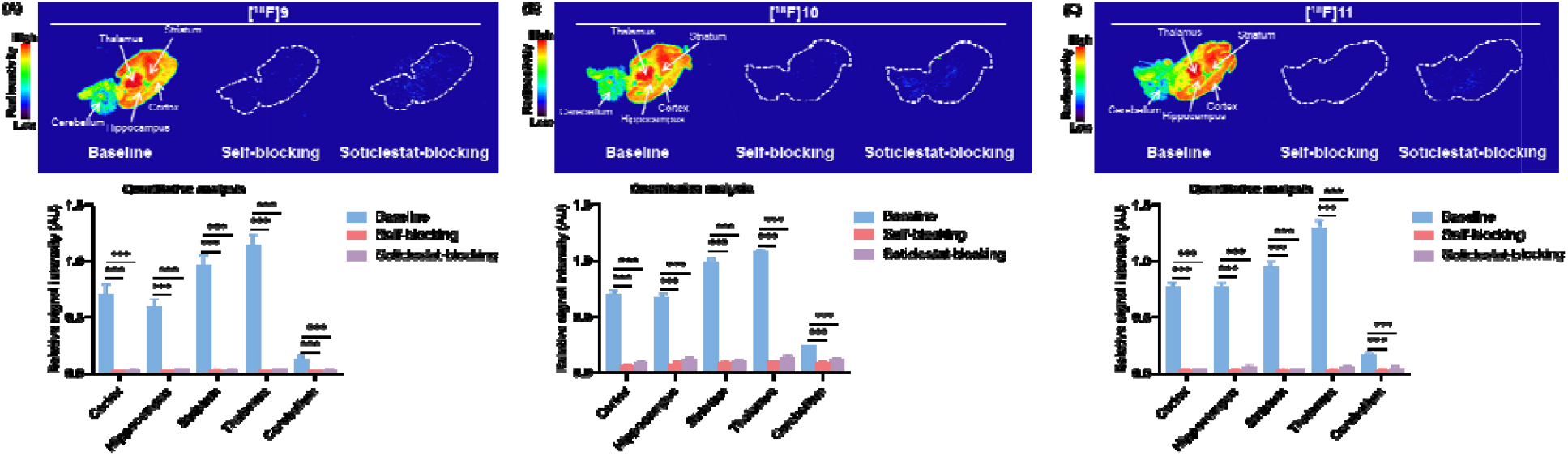
Representative *in vitro* autoradiograph analysis of (A) [^18^F]**9**, (B) [^18^F]**10**, and (C) [^18^F]**11** under baseline and blocking conditions (blocker = 10 μM). Asterisks (*) indicate statistical significance (\*\**p* ≤ 0.01, \*\*\**p* ≤ 0.001). Values represent mean ± SD, n = 3.

## CONCLUSION

In this study, we report the rational design and evaluation of three novel CYP46A1 PET tracers derived from the [^18^F]CHL-2205 scaffold through systematic modification of the left-side cyclopropyl moiety. Introduction of bridged, spirocyclic, and fused bicyclic motifs was well tolerated, yielding compounds **9**–**11** with subnanomolar CYP46A1 affinity comparable to CHL-2205. Efficient copper-mediated radiosynthesis afforded [^18^F] **9**–**11** with high radiochemical purity, molar activity, and excellent in vitro stability. *In vitro* autoradiography demonstrated regionally selective and specific binding of all tracers in rat brain sections, consistent with known CYP46A1 expression and comparable to [^18^F]CHL-2205. These results validate a scaffold-driven strategy for CYP46A1 PET tracer development and support further *in vivo* PET imaging studies to assess brain pharmacokinetics and translational potential.

## EXPERIMENTAL SECTION

### Materials and Methods

All reagents and solvents were purchased from commercial suppliers and used as received without further purification. Compounds **9**–**11** and their corresponding precursors **19**–**21** were synthesized following previously reported procedures.^26, 27^

### Radiochemistry

The aqueous [^18^F]fluoride was trapped on a Sep-Pak QMA Plus Light cartridge (Waters, cat. no. 186004540) and eluted in the reverse direction with a solution of TEAB (1 mg) in MeOH (1.0 mL). The eluate was azeotropically dried at 110 °C under a nitrogen stream with the addition of anhydrous MeCN (1.0 mL). A solution of the boronic ester precursors (**19**–**21**, 2 mg) and [Cu(Py)_4_OTf_2_] (8 mg) in dry DMAC/n-BuOH (200/100 μL) was then added, and the reaction mixture was heated at 110 °C for 15 min with the vial left uncapped. After completion, the crude reaction mixture was diluted with HPLC mobile phase and purified by semi-preparative HPLC on a Phenomenex Luna C18(2) column (5 μm, 10 × 250 mm), using MeCN/H_2_O (40/60, v/v, containing 0.1% NEt_3_) as the eluent.

### *In Vitro* Autoradiography

*In vitro* autoradiography was conducted using 20 μm rat brain cryosections embedded in Tissue-Tek® O.C.T. and stored at -80 °C. Prior to incubation, sections were pre-equilibrated for 10 min at room temperature in buffer 1 (50 mM Tris, 0.1% BSA). The sections were then incubated for 30 min in buffer 1 containing [^18^F]**9**–**11** (1 µci/mL). For blocking experiments, 10 μM of the corresponding unlabeled compounds **9**–**11** or the selective CYP46A1 inhibitor soticlestat was added during incubation. Following incubation, sections were washed in buffer 1 (1 × 5 min) and buffer 2 (50 mM Tris without BSA; 2 × 2 min), briefly rinsed twice in distilled water (5s each), and air-dried. The dried sections were exposed to a phosphor imaging plate (BAS-MS2025, GE Healthcare) for 12 h and subsequently scanned using an Amersham Typhoon imaging system (Cytiva, USA).

### Safety

No unexpected or unusually high safety hazards were encountered.

